# Reactivity-based proteomics uncovers ATP-induced proteome restructuring

**DOI:** 10.1101/2025.04.28.651067

**Authors:** Brian Críostóir Mooney, Farnusch Kaschani, Joy Chenqu Lyu, Markus Kaiser, Renier A. L. van der Hoorn

## Abstract

Chemical probes that label surface-exposed amino acid residues in native proteomes serve as powerful tools for studying native reactivities, yet their use has primarily been limited to drug discovery. In this study, we expand the application of reactivity proteomics by monitoring ATP-induced changes over 21,000 reactive lysine, serine, threonine, and tyrosine residues across native Arabidopsis proteomes using biotinylated N-hydroxysuccinimide (BioNHS) ester probes. ATP caused significant differential labelling. Labelling was reduced in ATP-binding pockets, consistent with ligand occupancy, whereas many sites outside ATP-binding regions showed either reduced or increased labelling in response to ATP. Structural modelling revealed that these differentially labelled sites are associated with ATP-induced conformational changes, altering lysine exposure. Notable examples of ATP-driven structural shifts include adenylate kinase ADK4, acyl-activating enzyme AAE3, cell division control protein CDC48A and its interacting partner PUX1, and ATP-induced 26S proteasome assembly. Furthermore, we show that reactivity proteomics using phosphonic acid-NHS (PhoNHS) followed by IMAC enrichment offers a complementary and expanded list of labelling sites. Our findings establish reactivity proteomics as a versatile and powerful platform for exploring protein conformational dynamics, metabolite–protein interactions, and regulatory mechanisms directly in native proteomes.

## Introduction

Reactivity-based proteomics is a chemical proteomics approach that provides insights into the reactivity and surface exposure of amino acid residues across whole proteomes (Hacker *et al*., 2017; Ward *et al*. 2017). These sites are monitored using small molecule probes that selectively label specific amino acid side chains. For example, *N-*hydroxysuccinimide (NHS) and sulfotetrafluorophenyl (STP) esters display strong proteomic reactivity predominantly by interacting with the primary amino group in the ε-amino side chain of lysine (Ward *et al*., 2017). Due to its hydrophilic nature, lysine is a common residue on protein surfaces making it a particularly suitable target for reactivity proteomics (Matos *et al*., 2018; Zanon *et al*., 2025).

The application of reactivity proteomics has so far been limited to biomedical science to identify ligandable hotspots within proteins that could be targeted by novel small molecule therapeutic drugs (Hacker *et al*., 2017; Ward *et al*. 2017). However, its potential to address wider questions in biology, including in plant science, remains largely untapped. We hypothesized that reactivity proteomics could be deployed as a discovery tool to characterise plant responses to various biological stimuli by monitoring proteome-wide changes in surface-exposure and reactivity at labelled sites. Dynamic responses including conformational changes, protein-protein interactions, ligand binding and post-translational modifications should theoretically result in labelling differentials on affected proteins **(Figure 1)**. Reactivity proteomics could be used in this manner to identify novel proteins and amino acid sites involved in the response to virtually any experimental treatment, including biotic or abiotic stressors.

**Figure 1.**
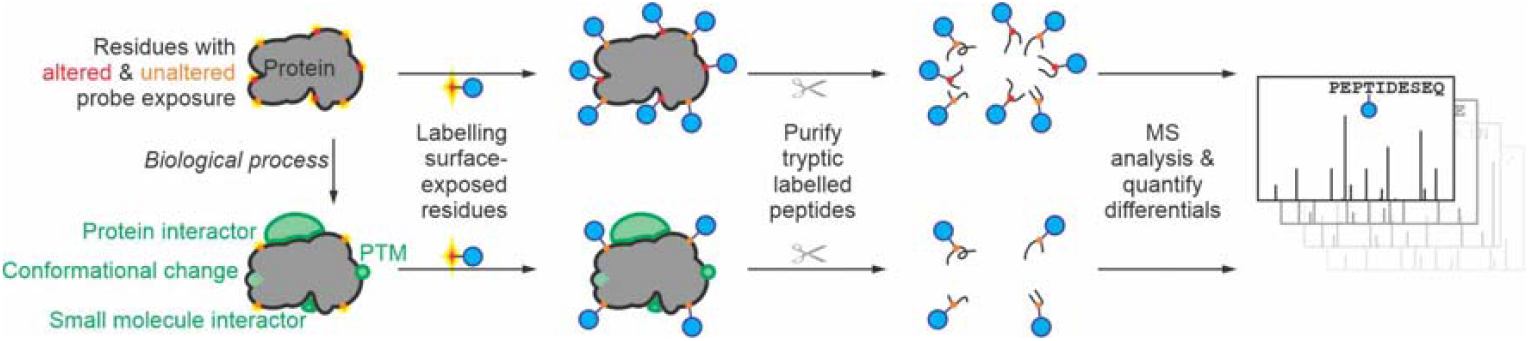
Concept of reactivity proteomics detecting dynamic proteomes. Reactivity probes (blue) label surface residues of native proteomes and can display differential labelling of sites that are altered by interactions with other proteins or small molecules, conformational change, and post-translational modifications (PTMs) (green).

The reactive moieties (or ‘warheads’) in reactivity probes are typically conjugated to affinity tags to facilitate enrichment and identification of the labelled sites by liquid chromatography tandem mass spectrometry (LC-MS/MS) (Morimoto & van der Hoorn, 2016). Biotin is the most commonly used reporter tag because its high affinity for (strept)avidin enables efficient purification of labelled peptide. However, stringent conditions required for elution often modifies eluted peptides and hinder their detection by MS (Kleinplenning *et al*., 2020). More recently, phosphonate-tagged probes have emerged as practical alternatives for protein labelling as the resulting phosphopeptides can be enriched with high sensitivity using immobilized metal affinity chromatography (IMAC) resins (Paulo *et al*., 2018; Kleinpenning *et al*., 2020; van Bergen *et al*., 2023).

Here, we establish reactivity proteomics in plants by monitoring reactive sites in the native proteome of *Arabidopsis thaliana* using a biotin-NHS (BioNHS) probe and enrichment of labelled peptides using avidin beads. We demonstrate the utility of reactivity proteomics to address biological questions by investigating differential labelling induced by ATP, revealing ATP binding sites and ATP-induced conformational changes. Finally, we describe an alternative reactivity proteomic strategy based on IMAC-enrichable phosphonic acid-NHS (PhoNHS) probe and compare this to BioNHS labelling. Together, these strategies facilitate the monitoring of over 21,000 unique sites across the native Arabidopsis proteome.

## Results

### Reactivity proteomics of plant proteomes

To explore plant proteomes with reactivity probes, we took advantage of a commercially available, amine-reactive biotin-NHS (BioNHS) probe (**Figure 2A**), and generated total proteomes from Arabidopsis cell cultures. We incubated these plant cell culture extracts with various BioNHS concentrations and detected biotinylated proteins on protein blots using a streptavidin-horseradish peroxidase conjugate. We detected strong proteome biotinylation that increases with increasing BioNHS concentrations (**Figure 2B**).

**Figure 2.**
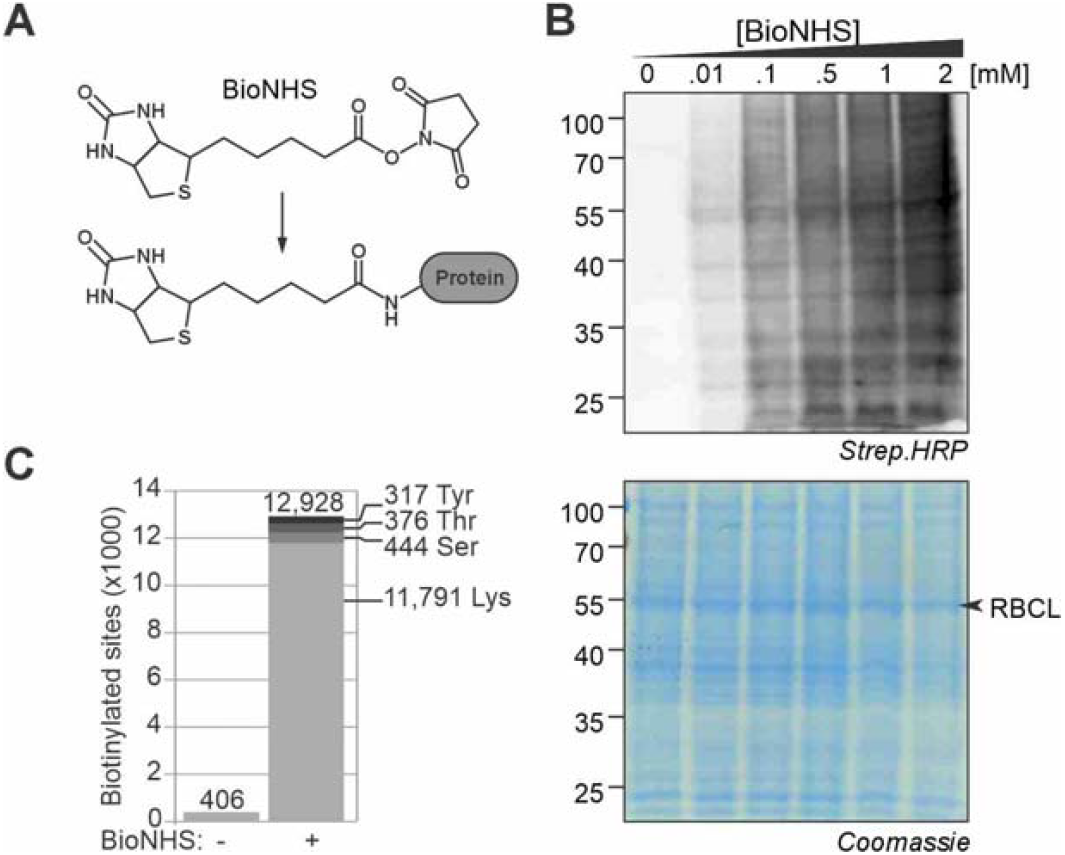
BioNHS labelling provides broad coverage of the Arabidopsis proteome. **(A)** Structure of BioNHS and its conjugation to a Lys residue. **(B)** BioNHS labels the Arabidopsis proteome in a concentration-dependent manner. Biotinylation was detected by streptavidin-HRP immunoblot. Total proteins were visualised using InstantBlue® Coomassie stain. **(C)** The number of unique KSTY sites biotinylated in the Arabidopsis proteome following BioNHS labelling or a no-probe control treatment.

To identify the labelling sites, Arabidopsis proteomes incubated with and without 1 mM BioNHS were digested with trypsin/LysC and biotinylated peptides were enriched on monomeric avidin agarose beads. NHS esters predominantly react with the primary amino group in the ε-amino side chain of lysines (K) but also with the hydroxyl groups of serines (S), threonines (T) and tyrosines (Y) (Ward *et al*., 2017). We therefore annotated the MS spectra with biotin modifications on K, S, T and Y residues. A total of 13,334 unique biotinylated KSTY sites in 3,503 proteins were identified (**Figure 2C; Supplementary Table S1**). 12,928 (97%) of these sites were biotinylated only upon incubation with BioNHS (**Figure 2C; Supplementary Table S1)**. Lysine was by far the most commonly labelled residue with 11,791 biotinylated lysine residues identified, accounting for over 91% of total labelling **(Supplementary Table S1)**. The remaining BioNHS-labelled sites comprised 444 serine (3.4%), 376 threonine (2.9%) and 317 tyrosine (2.5%) residues **(Supplementary Table S1)**. Biotinylation in the no-probe control samples **(Figure 2C, Supplementary Table S1)** includes the previously identified biotin-binding enzyme methylcrotonyl-CoA carboxylase subunit alpha (MCCA) at position K699, the predicted N6-biotinyllysine site (Baumgartner *et al*., 2001; Che *et al*., 2002).

### Reactivity-based proteomics detects ATP binding sites

To test whether reactivity proteomics could be used monitor alterations in a proteome induced by a cofactor, we pre-incubated the Arabidopsis proteome with or without 1 mM ATP and then labelled the proteome with 1 mM BioNHS (**Figure 3A**). We hypothesized that ATP would affect labelling not only by directly competing in ATP binding sites but also by ATP-induced protein-protein interactions and conformational changes.

**Figure 3.**
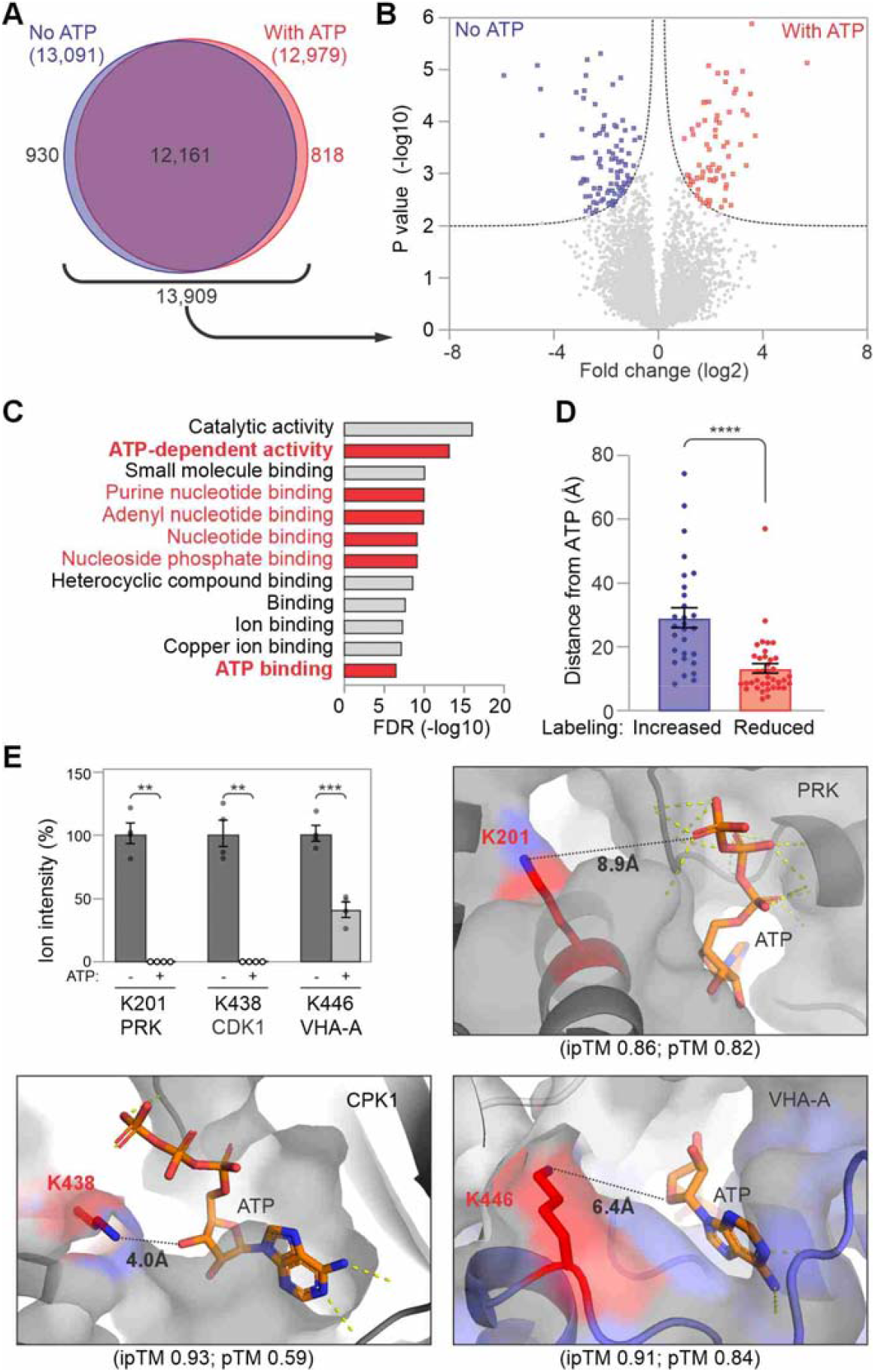
ATP restructures the reactive proteome. **(A)** Venn diagram showing overlap of labelled sites identified in the ATP-treated and untreated groups. Number of unique labelled KSTY sites in the ATP-treated and untreated samples (minimum 1 detection from 4 replicates). **(B)** Differentially labelled peptides following ATP treatment (minimum 3/4 detections in at least 1 treatment group). Right (blue) indicates sites with significantly reduced labelling in the presence of ATP. Left (red) indicates sites with significantly increased labelling in the presence of ATP. **(C)** Residues with altered labelling upon ATP treatment derived from proteins enriched in ATP-related GO terms (categories with ten lowest FDR values shown). **(D)** The proximity of residues with ATP-induced differential labelling to AlphaFold 3 predicted ATP-binding sites. Sites with reduced labelling are significantly closer to predicted ATP binding (Welch’s t-test, p < 0.0001). Error bars represent standard error. **(E)** ATP suppresses labelling at nearby residues. Three examples of ATP binding proteins that show suppressed labelling in the ATP binding pocket in the presence of ATP. Shown are AlphaFold3-predicted models with ATP and the shortest distance of ATP to the labelled lysine in PhosphoRibuloKinase (PRK), Calcium-dependent Protein Kinase 1 (CPK1), and V-type proton ATPase catalytic subunit A (VHA-A). **, P value Welsh t-test <0.01; ***, <0.001.

In this experiment, we identified 13,909 KSTY labelling sites in 3,255 proteins covering a wide range of GO terms (**Supplementary Table S2)**. Comparison between the ATP-treated and untreated samples revealed 166 sites in 137 proteins that were labelled differentially upon ATP pretreatment (**Figure 3B; Supplementary Table S3)**. These 137 proteins are significantly enriched for ATP-related functions, with ATP-dependent activity (GO:0140657), ATP binding (GO:0005524) and nucleotide binding (various GO terms) featuring among the most significantly enriched gene ontology (GO) terms (**Figure 3C; Supplementary Table S4**). This enrichment for ATP-related functions was not observed when the analysis was performed using the entire dataset of 3,255 labelled proteins (**Supplementary Table S5**).

Of the 137 proteins differentially labelled upon ATP treatment, 50 are known to bind ATP directly (**Supplementary Table S3**). To investigate whether ATP suppressed labelling near ATP binding sites, we used AlphaFold 3 (Abramson et al., 2024) to predict the structures of these proteins in complex with ATP. ATP treatment reduced labelling at 70 sites including 38 within ATP-binding proteins such as Calcium-dependent Protein Kinase 1 (CPK1), PhosphoRibuloKinase (PRK) and V-type proton ATPase catalytic subunit A (VHA-A) (**Figure 3E**). The affected sites within the set of ATP-binding proteins were on average less than 14 Å from the bound ATP ligand **(Figure 3D; Supplementary Table S6)**. Considering the estimated length of BioNHS (∼20 Å), it is likely that ATP binding obstructed BioNHS binding, causing reduced labelling at these sites.

In contrast, 96 sites in 84 proteins exhibited significantly *increased* labelling in response to ATP, including 28 sites from 22 ATP-binding proteins **(Supplementary Table S7)**. Within these 22 proteins, residues with increased labelling are significantly further from the predicted ATP-binding sites, at an average distance of 29 Å **(Figure 3D; Supplementary Table S7)**. In these cases, ATP binding may trigger conformational changes that lead to increased exposure of these residues. In addition to differential labelling of 49 known ATP-binding proteins, we also observed ATP-induced differential labelling in 88 proteins with no apparent ATP-binding function **(Supplementary Table S3)**. These differences may be caused by altered interactions with ATP binding proteins. We highlight five example proteins where differential labelling is caused by ATP-induced conformational changes.

### Case-1: ATP-suppressed labelling of acyl-activating enzyme AAE3

One of the differentially labelled proteins is the oxalyl-CoA synthetase ACYL-ACTIVATING ENZYME (AAE3), which is an ATP-binding protein for which crystal structures with and without ATP have been generated (PDB: 5EI0 and 5IE2, Fan *et al*., 2016). We detected 17 BioNHS-labelled KSTY sites in AAE3 (**Figure 4A**) but ATP only significantly suppressed labelling at three positions: K178, K411 and K482 (**Figure 4B**). The other 15 labelled sites were not suppressed by ATP pretreatment. Analysis of the crystal structure confirmed that two of these residues are in ATP-binding pockets (**Figure 4C**), located within 6.6 Å and 8.4 Å from ATP. ATP also suppressed labelling at K482, which is 16.9 Å of ATP, clearly outside the ATP binding pocket. ATP binding does not significantly change the structure of the region containing K482, nor does it change its exposure (Fan et al., 2016). This indicates that K482 reactivity reports on the presence of other ATP-induced interactions with AAE3.

**Figure 4.**
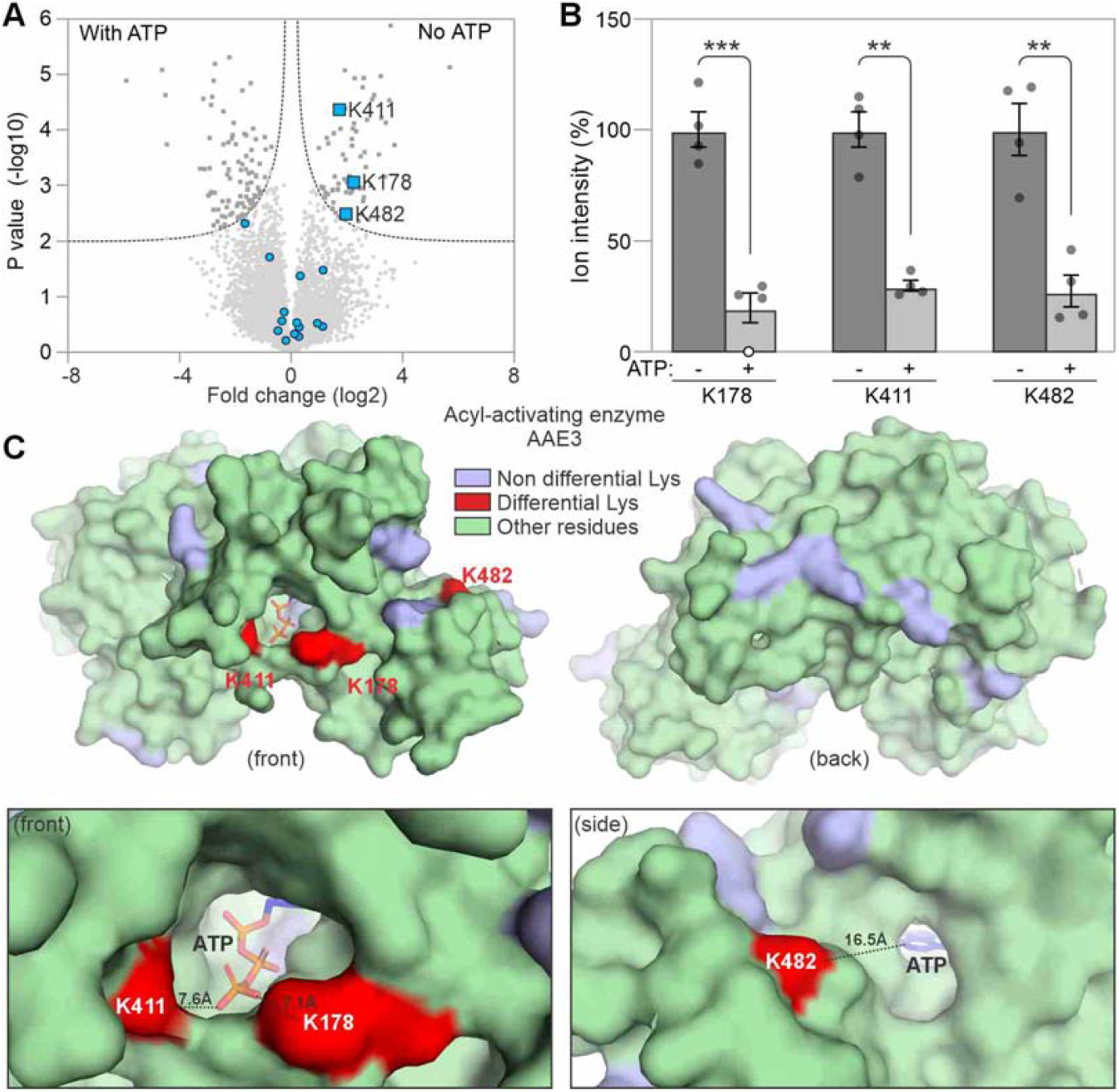
ATP supresses labelling in acyl-activating enzyme AAE3 inside and outside ATP pocket. **(A)** Volcano plot showing differentially labelled peptides following ATP treatment (minimum 3/4 detections in at least 1 treatment group). Highlighted are differentials (dark grey squares), non-significant labelling sites (light grey dots), and labelled sites in AAE3 (blue). **(B)** Intensity of differential BioNHS labelling in AAE3 in untreated and ATP-treated samples. Bars indicate means of 4 replicates +/-SEM expressed relative to the average untreated value. **, P value Welsh t-test <0.01; ***, <0.001. **(C)** Structure of AAE3 in complex with ATP (PDB: 5IE2, Fan *et al*., 2016). Highlighted are BioNHS-labelled residues (blue) that show reduced labelling in the presence of ATP (red). **(D-E)** K178, K411 and K482 surround the ATP binding site. Gold = sites with reduced labelling in the presence of ATP, red/orange = ATP.

### Case-2: ATP-induced and suppressed labelling of adenylate kinase ADK4

We detected 11 labelling sites in adenylate kinase ADK4 but ATP caused differential labelling on only three of these sites. Interestingly, labelling of K46 of ADK4 is reduced by ATP, whilst labelling at K81 and K222 in ADK4 is increased by ATP **(Figure 5A and 5B, Supplementary Table S3)**. AlphaFold3 predictions of ADK4 in complex with ATP indicate that K46 is within the ATP binding pocket (3.7 Å), explaining why K46 labelling is suppressed by ATP (**Figure 5C, Supplementary Table S7-S8)**. However, K81 and K222 are more distant from ATP (25.9 and 23.8, respectively) (**Figure 5C, Supplementary Table S7-S8)**. The increased labelling of K81 and K222 by ATP indicates that these two residues become more exposed upon ATP binding. Indeed, AlphaFold3 predictions of ADK4 with and without ATP indicate an ATP-induced conformational change (**Figure 5C**), consistent with previous observations that adenylate kinases undergo an ATP-induced conformational change from an open to a closed conformation (Pontiggia *et al*., 2008). However, the solvent exposure of K81 and K222 is not different with ATP, indicating that these residues report on altered interactions at this interface.

**Figure 5.**
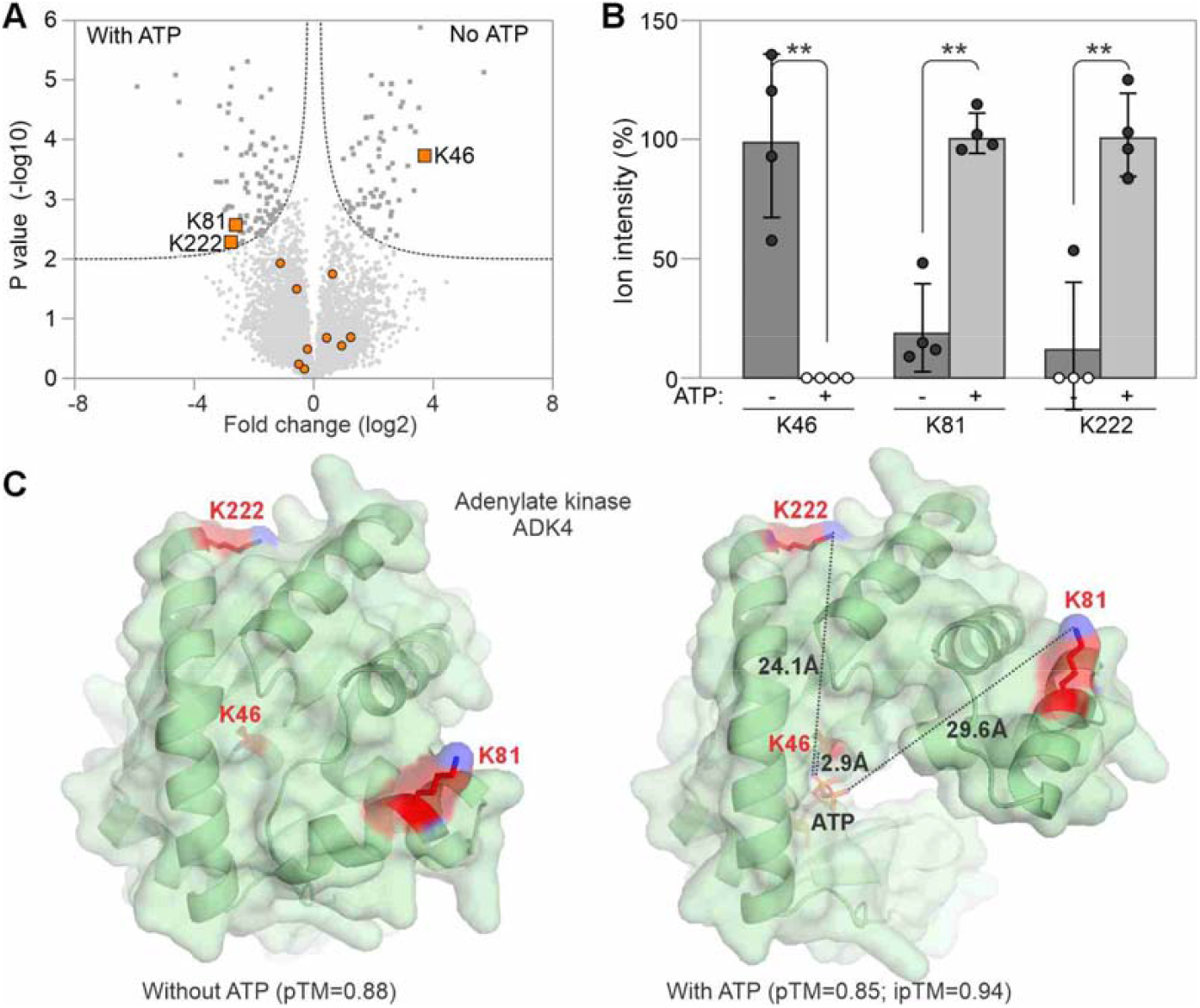
ATP suppresses and promotes reactivity labelling in adenylate kinase ADK4. **(A)** Volcano plot showing differentially labelled peptides following ATP treatment (minimum 3/4 detections in at least 1 treatment group). Highlighted are significant differentials (dark grey squares), non-significant labelling sites (light grey dots), and labelled sites in AAE3 (orange). **(B)** Intensity of differential BioNHS labelling in AAE3 in untreated and ATP-treated samples. Bars indicate means of 4 replicates +/-SEM expressed relative to the average untreated value. Asterisks denote statistical significance (FDR-adjusted p ≤ 0.05). **, P value Welsh t-test <0.01. **(C)** AlphaFold3-predicted structures of ADK4 with and without ATP. K46 locates in the ATP-binding pocket, whereas K81 and K222 are located distantly.

### Case-3: ATP-induced and suppressed labelling in ATPase CDC48A

We detected 18 labelling sites in CDC48A, a Cell Division Control protein that is a member of the family of AAA ATPases but only two of these sites are differentially labelled upon ATP treatment (**Figure 6A**). Interestingly, whilst labelling K239 is suppressed by ATP, labelling of K392 is significantly induced upon ATP (**Figure 6B)**. The ATP-suppressed labelling of K239 makes sense because this residue is proximate to the ATP-binding site K254 within the Walker A motif of the D1 domain (Aker *et al*., 2007), as indicated in the AlphaFold3-predicted structure (**Figure 6C**). K392, however, locates 16.7 Å from ATP and resides in a side cleft. AlphaFold3 predictions of the CDC48A hexamer with and without ATP indicates that this cleft is larger in the presence of ATP (**Figure 6D**), consistent with the increased labelling of this site, indicating that differential labelling of K392 reports a conformational change at the interface between CDC48A subunits.

**Figure 6.**
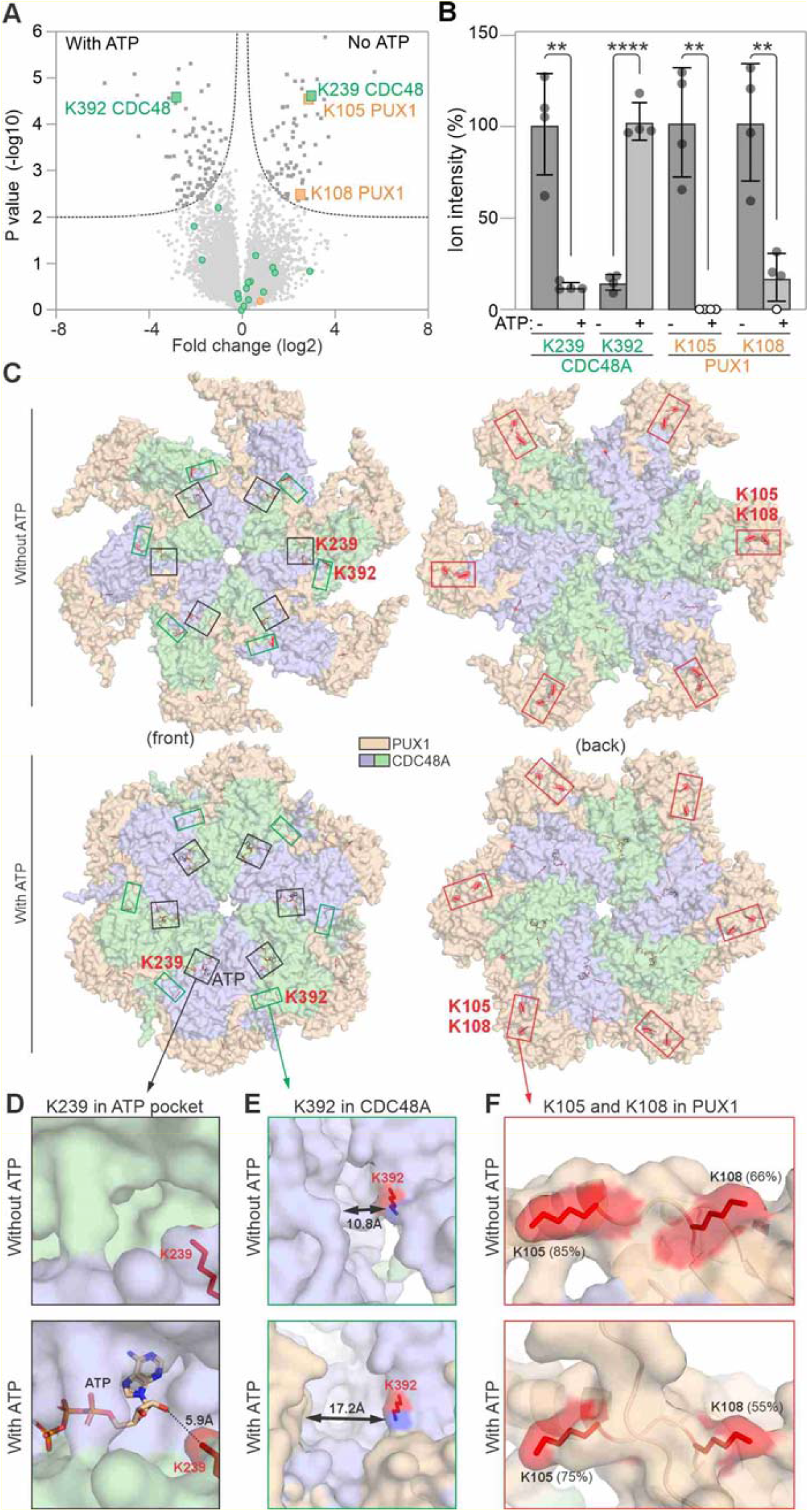
ATP triggers altered reactivity in CDC48A and CDC48-associated PUX1. **(A)** Volcano plot showing differentially labelled peptides following ATP treatment (minimum 3/4 detections in at least 1 treatment group). Highlighted are significant differentials (dark grey squares), non-significant labelling sites (light grey dots), and labelled sites in PUX1 (red) and CDC48A (green). **(B)** Intensity of differential BioNHS labelling in PUX1 and CDC48A in untreated and ATP-treated samples. Bars indicate means of 4 replicates +/-SEM expressed relative to the average untreated value. Asterisks denote statistical significance (FDR-adjusted p ≤ 0.05). **(C)** AlphaFold3 structural predictions of the CDC48A-PUX1 hexameric complex in the absence (top) and presence (bottom) of ATP. **(D)** Close-up of K239 in ATP binding pocket of CDC48A. **(E)** Close-up of K383 in CDC48A. **(F)** Close-up of K105 and K108 in PUX1 without and with ATP with percentage exposures calculated with the SASA function in PyMOL.

### Case-4: ATP-induced labelling of CDC48A-interacting PUX1

We also detected ATP-reduced labelling in UBX domain-containing protein 1 (PUX1) at K105 and K108 (**Figure 6A**). PUX1 is not known to bind to ATP directly, but these two residues are located within the UBX domain, which interacts with AAA+ ATPase CDC48A (Rancour *et al*., 2004; Park *et al*., 2007; Zhang *et al*., 2021). Structural predictions of PUX1-CDC48A heterohexamers with and without ATP using AlphaFold3 indicated that K105 and K108 are partly obstructed in the ATP-bound conformation by the altered position of the C-terminal portion of PUX1, resulting in a reduced solvent-accessible surface area (SASA) **(Figure 6C)**. These data are consistent with previous reports that PUX1 negatively regulates CDC48A activity in the presence of ATP by disturbing its active hexameric form (Park *et al*., 2007).

### Case-5: ATP-suppressed labelling of α2 proteasome subunit

Besides PUX1, ATP also reduced labelling at K176 in the α2 subunit of the proteasome (PAB1, **Figure 7A**), even though this protein does not bind ATP itself. PAB1 is a component of the two outer α-rings of the 20S core particle (CP) of the proteasome, which also contains two inner rings, each of seven β subunits, including the catalytic subunits. K176 is fully exposed at the CP in the Cryo-EM structure of the 20S proteasome of spinach (**Figure 7C**), explaining its labelling in the absence of ATP (7QVE, Kandolff et al., 2022, **Figure 7B**). In the presence of ATP, however, the CP associates with the regulatory particle (RP) to assemble into the 26S proteosome (Liu et al., 2006). Cryo-EM structures of the 26S proteasome (6EPC, Guo et al., 2018) demonstrate that K176 is covered by the RPN6 (At1g29150) subunit of the RP (**Figure 7D**). Thus, reduced labelling of K176 in PAB1 is caused by an ATP-induced protein-protein interaction.

**Figure 7.**
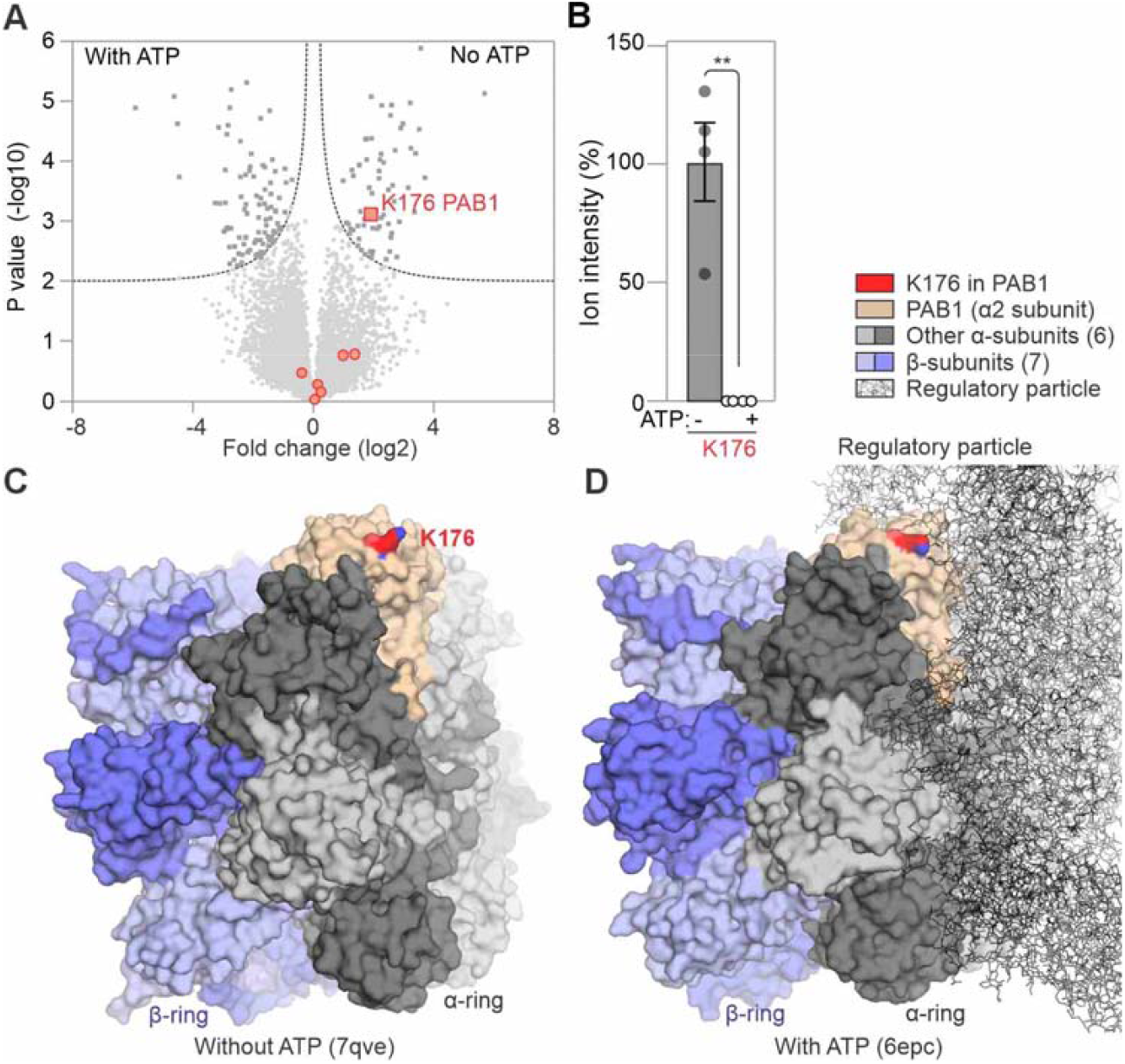
ATP-induced 26S proteasome assembly suppresses K176 labelling in α2 subunit. **(A)** Volcano plot showing differentially labelled peptides following ATP treatment (minimum 3/4 detections in at least 1 treatment group). Highlighted are significant differentials (dark grey squares), non-significant labelling sites (light grey dots), and labelled sites in PAB1 (red). **(B)** Intensity of differential BioNHS labelling on K176 of PAB1 in untreated and ATP-treated samples. Bars indicate means of 4 replicates +/-SEM expressed relative to the average untreated value. Asterisks denote statistical significance (FDR-adjusted p ≤ 0.05). **(C)** Cryo-EM Structure of the 20S core particle of the spinach proteasome (7qve, Kandolff et al., 2022). Only two of the four rings are shown: the β-ring containing the three catalytic subunits (blue surface) and the α-ring (grey surface) with the α2 (PAB1, light red surface) with the K176 (red surface) highlighted. **(D)** Cryo-EM structure of the ground state rat 26S proteasome (6epc, Guo et al., 2018) containing the regulatory particle (black lines) docked on the core particle (blue/grey surface), hiding K176 (red surface) in the α2 subunit (light red surface).

Besides these cases, **Supplementary Table S3** lists many additional ATP-dependent differential labelling sites that are not associated with direct ATP binding, but warrant further investigation. Labelling at K18 and K136 in ascorbate peroxidase APX1, for instance, is reduced and increased in the ATP sample, respectively. Labelling of T163 in glutathione transferase GSTF6 is strongly decreased upon ATP binding, whereas S159 labelling in GSTF6 is strongly increased upon ATP binding. Differential labelling of metabolic enzymes also indicates ATP-induced regulation: labelling of K348 in phosphoenolpyruvate carboxylase PEPC and K240 in malate dehydrogenase MDH2 are both increased upon ATP treatment, whilst labelling of K277 in cytosolic fructose biphosphatase CYFBP and K53 in chloroplastic ferredoxin-nitrate reductase NIR1 are suppressed by ATP. How ATP induces differential labelling in these proteins requires further investigation.

### PhoNHS labels a distinct set of reactive sites in the plant proteome

Finally, we investigated the use of an alternative phosphonate-IMAC enrichment strategy. We custom-synthesized a phosphonic acid-NHS (PhoNHS) probe (**Figure 8A**) and tested its reactivity with the Arabidopsis proteome. Robust labelling was detected at concentrations ≥100 μM PhoNHS using the ProQ diamond phosphoprotein gel stain (**Figure 8B**).

**Figure 8.**
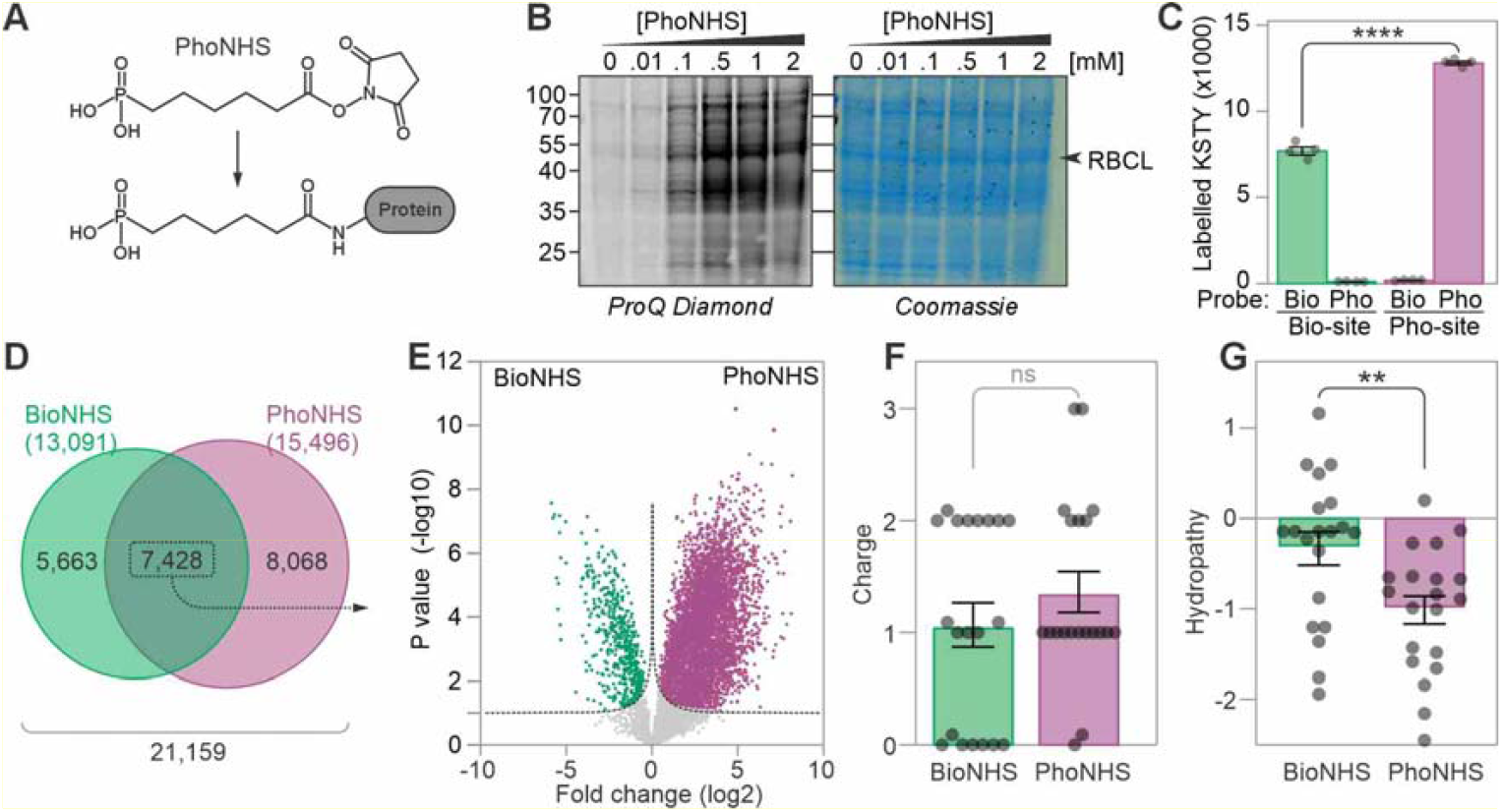
Expanded reactivity profiling with PhoNHS probes. **(A)** Chemical structure of PhoNHS and its conjugate on Lys residues. **(B)** BioNHS labels the Arabidopsis proteome in a concentration-dependent manner. Biotinylation was detected by streptavidin-HRP immunoblot. Total proteins were visualised using InstantBlue® Coomassie stain. **(C)** Number of KSTY residues labelled with BioNHS and PhoNHS detected each replicate. Arabidopsis proteome was labelled with BioNHS and purified on avidin beads (Bio) or labelled with PhoNHS and purified by IMAC (Pho) in four replicates each. Biotinylated and Phosphorylated KSTY sites were detected in both enrichment procedures. Error bars represent standard error of n=4 replicates. ****, P value Students t-test <0.0001. **(D)** Venn diagram comparing the total KSTY sites labelled by PhoNHS and BioNHS. **(E)** Volcano plot comparing the intensity of sites labelled by both BioNHS and PhoNHS. Left: stronger intensities upon BioNHS labelling; right: stronger intensities upon PhoNHS labelling. FDR = 0.01. **(F)** Net charge of the 20 sites most differentially labelled by PhoNHS or BioNHS determined using the GRAVY index. P value, Welsh t-test non-significant (ns). **(F)** Hydropathy scores of the 20 most differentially labelled sites using either PhoNHS or BioNHS. **, P value Welsh t-test <0.01.

We next compared labelling by 1 mM PhoNHS with 1 mM BioNHS. We identified 66% more KSTY sites labelled with PhoNHS than with BioNHS (**Figure 3C**). The 15,496 unique PhoNHS sites included 14,320 lysine (∼ 92%), 626 serine (∼ 4%), 339 threonine (∼ 2%) and 211 tyrosine (∼ 1%) residues (**Supplementary Table S8)**. When compared with the 13,091 residues labelled by BioNHS in this experiment **(Figure 8C)**, only 7,428 sites were labelled with both probes, while 8,068 and 5,663 were exclusively labelled by either PhoNHS or BioNHS, respectively (**Figure 8C**), displaying remarkably distinct labelling. Combined, these two probes provide coverage of 21,159 unique KSTY sites in the detected proteome. This includes 18,812 unique lysines, representing 12.6% of the 149,307 lysines present in our dataset of 5,033 detected proteins **(Supplementary Table S9)**.

Of the 7,428 sites labelled by both probes, 4,946 (∼67%) were detected with a significantly higher intensity using the PhoNHS-IMAC approach (**Figure 8E**). To understand the basis of these differences in probe selectivity, we assessed the chemical properties of 7-amino acid long sequences including the 3 residues sequentially up- and downstream of the 20 most strongly differentially labelled sites **(Supplementary Table S10**). This analysis indicated that the charge of the adjacent residues has little effect (**Figure 8E**) but that PhoNHS preferentially labels sites surrounded by hydrophilic residues while BioNHS performs better when labelling sites adjacent to hydrophobic residues (**Figure 8F**). Taken together, these data demonstrate that PhoNHS labelling highlights a complementary and distinct set of KSTY sites in the proteome.

## Discussion

We demonstrated that reactivity profiling is a powerful platform to monitor proteome-wide changes in protein reactivities. Using lysine-reactive probes and an approach based on the enrichment of labelled peptides, we were able to monitor 13,000-21,000 reactive lysine, serine, threonine and tyrosine (KSTY) residues in the Arabidopsis proteome, significantly expanding the known landscape of reactive residues and providing a valuable resource for the plant science community. Notably, we showed that reactivity profiling can not only be used to monitor small molecule binding sites, exemplified by ATP, but it can also monitor structural changes in the proteome induced by small molecule binding. These changes include conformational changes, altered protein-protein interactions, post-translational modifications and small molecule interactions.

Notably, we also identified 406 biotinylated KSTY sites in our no-probe treatment control samples. This number far exceeds the handful of Arabidopsis proteins that are currently known to be biotinylated *in vivo*. Most of these labelled peptides, however, were found by matching the MS1 signal, not by MS itself, and are therefore likely false positives. Disabling the matching peaks will remove the signals in the no-probe control, but will also reduce the quantification in probe-labelled samples, though labelled peptides were identified in at least one of the labelled samples.

Following labelling with BioNHS, however, the number of biotinylated peptides increased dramatically to over 13,000. Most biotinylated peptides contain a biotinylated lysines but biotinylated serine, threonine and tyrosine residues were also detected. The 4,451 proteins detected in this experiment carry a total of 128,844 lysine, 169,500 serine, 101,453 threonine and 55,615 tyrosine residues, of which 9.15% lysine, 0.26% serine, 0.37% threonine and 0.57% tyrosine were detected as biotinylated residues. Many KSTY sites are not biotinylated because they are hidden in protein structures but also because some peptides are too small or too large to be detected. Please also note that lysine biotinylation would block trypsin processing, which is why several trypsin mis-cleavages were permitted in the searches.

We deployed reactivity proteomics to sensitively detect ATP binding and ATP-induced conformational changes. ATP treatment led to significant differences in BioNHS labelling in a set of proteins strongly enriched for ATP-related functions. In ATP-binding proteins like AAE3 and ADK4, most residues with ATP-reduced labelling were very close to the ATP binding pocket at an average distance of 13 Å, strongly suggesting their reduced accessibility for labelling following ATP binding. Reactivity proteomics could be used in this manner to monitor the availability of ligand binding sites. This strategy contrasts with the approach of using ligand-derived probes (*e*.*g*. acyl-ATP) to directly label binding sites (Patricelli *et al*., 2007; Villamor *et al*., 2013; Franks *et al*., 2019). Thus, reactivity proteomics is a highly flexible discovery tool that can be used to investigate the response to virtually any type of chemical treatment or biological stimulus without the need to synthesize custom ligand-based probes **(Figure 1)**.

In many cases, ATP treatment led to differential labelling at multiple sites, sometimes in opposing directions. In ADK4 for example, ATP reduced labelling at K46 in its ATP binding pocket, but led to increased labelling at K81 and K222, which reflects the previously described ATP-induced conformational change in ADK4 (Pontiggia *et al*., 2008). However, structural models suggest that ATP-induced increased reactivity of K81 and K222 is not caused by an increased solvent exposure from ADK4 itself, which implies that this surface interacts with proteins or other molecules in the absence of ATP. So far, no interactions that could explain this differential labelling have been described. Also CDC48A displayed ATP-suppressed labelling in its ATP binding pocket at K239, and ATP-induced labelling at K392. The latter is likely caused by the described ATP-induced conformational change of CDC48A (ref). Structural predictions of the CDC48A hexamer indeed indicated that K392 is exposed in a cleft that becomes larger when the hexamer contains ATP.

ATP also reduced labelling in proteins that are not known to bind to ATP. As canonical ATP binding motifs are also absent in these proteins, we consider it unlikely that they represent a novel set of ATP-binding proteins. For example, residues K105 and K108 of PUX1 exhibited decreased labelling in response to ATP, despite no predicted ATP-binding activity. However, these residues reside in a UBX domain that mediates an interaction with CDC48A, an AAA ATPase that undergoes ATP-induced conformational changes (Rancour *et al*., 2004; Park *et al*., 2007; Zhang *et al*., 2021; Huntington *et al*., 2025). These findings expand the applicability of reactivity proteomics beyond ligand-binding to the study of dynamic protein-protein interaction interfaces in response to biochemical cues.

Biotin-conjugated chemical probes such as BioNHS have been widely used for protein labelling because of the ability to enrich biotinylated proteins or peptides using streptavidin and related molecules. However, purification of biotinylated peptides has several limitations: i) Many proteins stick non-specifically to streptavidin beads; ii) biotin is a relatively large modification that shifts many labelled peptides out of the MS detection window; and (iii) because of the high affinity of the biotin-streptavidin interaction, the elution of biotinylated peptides requires harsh conditions, resulting in the modification of 95-99% of the biotinylated peptides, such that these peptides can no longer be recognized by MS analysis (Kleinpenning *et al*., 2020). Attempts to improve the detection of labelled peptides include the use of desthiobiotin and/or neutravidin to reduce the affinity, and the use of cleavable linkers in probes (Verhelst *et al*., 2007; Morimoto & van der Hoorn, 2016), but these methods are also often inefficient and still cause peptide modifications. As an alternative, phosphonate-conjugated labelling reagents generate phosphopeptides that can be enriched efficiently using immobilized metal-ion affinity chromatography (IMAC) resins (Ficarro *et al*., 2002; Kleinpenning *et al*., 2020). A distinct advantage of phosphonic acid versus biotin-based probes is that the phosphonate modification at labelled residues is distinct from the natural phosphorylation signature, reducing the confounding effects from endogenously modified peptides.

To expand the application of reactivity proteomics in plants, we developed a phosphonic acid-NHS (PhoNHS)-IMAC pipeline using a custom-synthesized probe. Using PhoNHS, we identified 15,459 labelled KSTY sites across the Arabidopsis proteome, including an additional 8,068 (∼52%) that were not labelled with BioNHS. When compared with PhoNHS, BioNHS preferentially labels sites in regions with higher hydrophobicity. PhoNHS meanwhile preferentially labels regions with comparably higher net charge, consistent with the fact that this probe negatively charged at physiological pH. A similar comparison of phosphonate versus biotin labelling performance by Kleinpenning *et al*. (2020) retrieved a higher proportion of non-specific enrichment when using a biotin-based approach, although in this case biotinylated proteins were enriched followed by on-bead digest, whereas labelled peptides were enriched in our study. These findings indicate while both BioNHS and PhoNHS provide broad proteome coverage, their suitability for specific experiments may vary based on their chemical characteristics. The complementary coverage of PhoNHS and BioNHS also highlights the potential for the use of diverse chemical probes to map the reactive proteome with even broader proteome-wide coverage.

The theoretical maximum number of labelling sites in proteomes is limited by several factors, including protein abundance, residue exposure and reactivity. The number of labelling sites that are ultimately identifiable is also affected by the sensitivity of the MS analysis. For example, in our experiment, labelling at sites within tryptic peptides that fall outside the MS detection window based on their size will not be captured. Reactivity proteomics following digest with alternative proteases (*e*.*g*. chymotrypsin) could therefore lead to the identification of additional labelling sites in Arabidopsis.

NHS probes can provide broad proteome coverage particularly because of the prevalence of lysines on protein surfaces (Matos *et al*., 2018). However, many other amino acids theoretically be targeted by reactivity probes. For example, iodoacetamide probes have been used to monitor cysteine reactivity in mice and the plant pathogen *Pseudomonas syringae* (Xiao *et al*., 2020; Morimoto *et al*., 2022). Considering that the probes used here enabled monitoring of over 21,000 KSTY sites in Arabidopsis, we anticipate that the use of probe cocktails targeting diverse amino acids in parallel could provide unrivalled proteome coverage amounting to many tens of thousands of sites. Future work could also extend to the application of reactivity proteomics *in vivo* in plants, including via the direct infiltration of probes into plant tissues.

In conclusion, these findings establish reactivity proteomics as a novel and versatile tool for plant proteomics, capable of detecting both small molecule binding and secondary-effect conformational changes in complex proteomes. This approach opens new possibilities for investigating protein function in plants.

### Experimental procedures

#### Arabidopsis cell culture

Cells were obtained from a suspension culture of wild-type *Arabidopsis thaliana* (L.) Heynh. ecotype Landsberg erecta, derived from an original stock established by May and Leaver (1993). The culture was maintained in the light at 22 °C on a rotary shaker in Murashige and Skoog (MS) medium supplemented with 160 mM glucose, 0.5 mg/L naphthalene acetic acid and 0.05 mg/L kinetin. Cells were harvested for experiments 5-7 days after subculturing.

#### Protein extraction

Arabidopsis cell samples were collected by centrifugation of 1 mL aliquots of the cell suspension. Cell pellets were flash frozen in liquid nitrogen and stored at -80 °C. Cells were lysed via addition of 800 μL of extraction buffer (50 mM HEPES pH 7, 1% PVPP, 5 mM MgCl2, 10% glycerol, 0.3% Triton-X). Lysates were incubated with the Benzonase® nuclease (25 units per mL) for 30 minutes on ice before centrifugation at 15,000 x g to remove nucleic acids and cellular debris. Soluble proteins in the supernatant were retained and combined into a single tube. Protein yields were measured using the DC protein assay (Bio-Rad).

#### ATP treatment

The Arabidopsis protein extract was divided into 8 identical aliquots of 700 μg protein each. 1 mM ATP or a mock control was added to 4 replicates each and samples were incubated at room temperature for 30 minutes before labelling with NHS reagents.

#### Labelling with NHS reagents

The biotin-NHS (BioNHS) probe used in this study is commercially available from Thermo Scientific™ as EZ-Link™ NHS-Biotin (20217). The phosphonic-acid NHS (PhoNHS) probe was synthesized by Haoyuan ChemExpress Co., Ltd. Stock solutions of NHS reagents were prepared in DMSO. Protein extracts were incubated with 1 mM BioNHS or 1 mM PhoNHS or an equivalent volume of DMSO (mock) on a rotary wheel for 1 hour at room temperature. BioNHS labelling was assessed by immunoblotting using Streptavidin-HRP (Sigma; S2438). PhoNHS labelling was determined using the Pro-Q™ Diamond Phosphoprotein Gel Stain (Invitrogen) following SDS-PAGE. To terminate labelling, excess NHS reagent was removed in the supernatant following acetone precipitation and centrifugation.

#### Tryptic digest

Protein pellets were resuspended in denaturing buffer (6 M urea, 2 M thiourea in 50 mM ammonium bicarbonate) and incubated for 10 minutes at room temperature. Proteins were reduced with 10 mM TCEP and incubated at room temperature in the dark for 30 minutes. For cysteine alkylation, 50 mM iodoacetamide was added and samples were incubated in the dark for a further 20 minutes. The urea concentration was reduced to approximately 1 M by the addition of 50 mM ammonium bicarbonate. Trypsin/Lys-C (Promega) was added to the samples at a 1:100 ratio of protease to protein. Digestion reactions were incubated overnight with shaking at 37 °C.

#### Peptide desalting

Peptides were desalted using Sep-Pak C18 cartridges (Waters) and stored at -20 °C. Briefly, tryptic peptides were acidified by the addition of trifluoroacetic acid (TFA) to 0.5%. The peptide samples were then diluted in solution A (2% acetonitrile, 0.1% TFA) before being passed through the C18 cartridge using a needleless syringe. After washing with solution A, desalted peptides were eluted from the C18 membrane with solution B (65% acetonitrile, 0.1% TFA). Purified peptides were dried in a vacuum concentrator.

### Enrichment of biotin-NHS labelled peptides

Desalted peptides were resuspended in 25 mM ammonium bicarbonate. Peptide concentration was measured using a BCA assay (Pierce Quantitative Colorimetric Peptide Assay). Biotinylated peptides were enriched using monomeric avidin agarose beads (Pierce). Beads were washed 3 times with H_2_O. 500 μg input peptides were added to the washed beads and incubated on a rotating wheel for 1 hour at room temperature. Beads were washed 3 times with 300 μL MS Grade H_2_O to remove nonspecific peptides. Biotinylated peptides were eluted from the beads with 500 μL elution buffer (80% acetonitrile; 1 % TFA) for 30 minutes at room temperature followed by 2 minutes at 96 °C. Beads were pelleted by centrifugation and eluted peptides in the supernatant were moved to a clean tube. Peptides were dried down in a vacuum concentrator.

### Enrichment of phosphonic acid-NHS labelled peptides

Labelled phosphopeptides were enriched using MagReSyn Zr-IMAC and Ti-IMAC HP magnetic microparticles (ReSyn Biosciences). Dried, desalted peptides were resuspended in 200 μL IMAC loading buffer (0.1 M glycolic acid in 80% acetonitrile, 5% TFA). Samples were centrifuged to pellet and remove insoluble material. Zr and Ti-IMAC beads were combined at a 1:1 ratio and equilibrated in loading buffer before the addition of the peptides. Peptides were incubated with the beads for 30 minutes at room temperature on a rotating wheel. Flow-through supernatant was removed using a magnetic separation rack. Beads were washed with loading buffer, wash buffer 1 (80% acetonitrile; 1% TFA) and wash buffer 2 (10% acetonitrile; 0.2% TFA) for 2 minutes each with gentle agitation. Bound peptides were eluted by incubation with 1% ammonium hydroxide for 20 minutes. 2.5% formic acid was added to the eluate. Peptides were dried down using a vacuum concentrator.

### Analysis of protein structures

AlphaFold structural models were produced using AlphaFold 3 (https://alphafoldserver.com) and visualised using PyMOL.

### Sample clean-up for LC-MS/MS

Peptides were desalted on home-made C18 StageTips containing two layers of an octadecyl silica membrane (CDS Analytical, Oxford, PA, USA). All centrifugation steps were carried out at room temperature. The StageTips were first activated and equilibrated by passing 50 μL of methanol (600 × g, 2 min), 80% (v/v) acetonitrile (ACN) with 0.5% (v/v) FA (600 × g, 2 min) and 0.5% (v/v) FA (600 × g, 2 min) over the tips. Next, the acidified tryptic digests were passed over the tips (800 × g, 3 min). The immobilized peptides were then washed with 50 μL and 25 μL 0.5% (v/v) FA (800 × g, 3 min). Bound peptides were eluted from the StageTips by application of two rounds of 25 μL 80% (v/v) ACN with 0.5% (v/v) FA (800 × g, 2 min). After elution from the StageTips, the peptide samples were dried using a vacuum concentrator (Eppendorf, Hamburg, Germany) and the peptides were dissolved in 15 μl 0.1% (v/v) FA prior to analysis by MS.

### LC-MS/MS Analysis

LC-MS/MS analysis of peptide samples were performed on an Orbitrap Fusion Lumos mass spectrometer (Thermo Scientific, Waltham, MA, USA) coupled to a Easy nLC 1200 or to a Vanquish Neo ultra high-performance liquid chromatography (UHPLC) system (Thermo Scientific, Waltham, MA, USA) that were operated in the one-column mode. The analytical column was a fused silica capillary (inner diameter 75 μm, outer diameter 360 μm, length 28 cm; New Objective, Littleton, MA, USA) with an integrated sintered frit packed in-house with Kinetex 1.7 μm XB-C18 core shell material (Phenomenex, Aschaffenburg, Germany). The analytical column was encased by a PRSO-V2 column oven (Sonation, Biberach, Germany) and attached to a nanospray flex ion source (Thermo Scientific, Waltham, MA, USA). The column oven temperature was set to 50 °C during sample loading and data acquisition. The LCs were equipped with two mobile phases: solvent A (2% ACN and 0.2% FA, in water) and solvent B (80% ACN and 0.2% FA, in water). All solvents were of UHPLC grade (Honeywell, Charlotte, NC, USA). Peptides were directly loaded onto the analytical column with a maximum flow rate that would not exceed the set pressure limit of 950 bar (usually around 0.5 – 0.6 μl min–1) and separated on the analytical column by running a 105 min gradient of solvent A and solvent B at a flow rate of 300 nL/min (start with 3% (v/v) B, gradient 3% to 9% (v/v) B for 6:30 min, gradient 9% to 30% (v/v) B for 62:30 min, gradient 30% to 50% (v/v) B for 24 min, gradient 50% to 100% (v/v) B for 2:30 min and 100% (v/v) B for 9:30 min). The mass spectrometer was controlled by the Orbitrap Fusion Lumos Tune Application and operated using the Xcalibur software. The MS settings for the different experiments are provided in Supplemental **Table S11**.

### Data processing and analysis

RAW spectra were submitted to an Andromeda (Cox et al., 2011) search in MaxQuant (2.0.3.0) using the default settings (Cox & Mann 2008). Label-free quantification and match-between-runs was activated (Cox et al., 2014). The MS/MS spectra data were searched against the Uniprot A. thaliana reference proteome UP000006548_3702_OPPG.fasta (one protein per gene; 27501 entries). All searches included a contaminants database search (as implemented in MaxQuant, 245 entries). The contaminants database contains known MS contaminants and was included to estimate the level of contamination. Andromeda searches allowed oxidation of methionine residues (16 Da, M) as variable modification. Depending on the experiment also modification of peptides by Bio-NHS (226 Da, specificity KSTY) or Phospho-NHS (178 Da, specificity KSTY) were allowed as variable modification. Carbamidomethylation on Cystein (57) was selected as static modification. Enzyme specificity was set to “Trypsin/P”. The instrument type in Andromeda searches was set to Orbitrap and the precursor mass tolerance was set to ±20 ppm (first search) and ±4.5 ppm (main search). The MS/MS match tolerance was set to ±0.5 Da. The peptide spectrum match FDR and the protein FDR were set to 0.01 (based on target-decoy approach). For protein quantification unique and razor peptides were allowed. Modified peptides were allowed for quantification. The minimum score for modified peptides was 40. Label-free protein quantification was switched on, and unique and razor peptides were considered for quantification with a minimum ratio count of 2. Retention times were recalibrated based on the built-in nonlinear time-rescaling algorithm. MS/MS identifications were transferred between LC-MS/MS runs with the “match between runs” option in which the maximal match time window was set to 0.7 min and the alignment time window set to 20 min. The quantification is based on the “value at maximum” of the extracted ion current. At least two quantitation events were required for a quantifiable protein. Further analysis and filtering of the results was done in Perseus v1.6.10.0. (Tyanova et al., 2016). Comparison of protein group quantities (relative quantification) between different MS runs is based solely on the LFQ’s as calculated by MaxQuant, MaxLFQ algorithm (Cox et al., 2014). Data were presented using GraphPad Prism. GO term enrichment analysis was performed at geneontology.org using the molecular function GO aspect. Venn diagrams were produced using BioVenn (Hulsen *et al*., 2008).

### Peptide property analysis

Differentially labelled sites from the bioNHS vs. phoNHS comparison were assessed for differences in hydropathy and net charge. Hydropathy was determined according to the grand average of hydropathy (GRAVY) index using www.gravy-calculator.de. Net charge was determined by uploading the 7-amino acid long sequences to www.pepcalc.com.

## Data availability

The mass spectrometry proteomics data for the on-bead digestions have been deposited to the ProteomeXchange Consortium via the PRIDE (Vizcaíno et al., 2016) partner repository (https://www.ebi.ac.uk/pride/archive/) with the dataset identifier PXD073071. During the review process the data can be accessed via a reviewer account (Username; Password: 0HYo3kqBcgn5)

## Funding

This project was financially supported by ERC-2020-AdG project 101019324 ‘ExtraImmune’ (BCM, RH), and BBSRC project DDT00230 (JL).

## Author contributions

RH conceived the work. BCM conducted experiments. FK generated proteomics data. BCM and RH wrote the manuscript and prepared figures with input from all co-authors.

## Conflicts of interest

none declared.

---

**Supplemental Table S11.** MS settings for whole proteome analysis.

| Project | General | MS1 | MS2 | Comments; special settings |
| --- | --- | --- | --- | --- |
| ACE_0717 | Tune v3.3.2782.28<br>Gradient: 105 min | Analyzer: FT<br>Res.: 240000<br>SR: 375 - 1500<br>AGC: Standard<br>AcT: Auto<br>RF: 30<br>SF: --<br>DDM: CT/3sec | Analyzer: FT<br>Res./ScR: 7500/-<br>SR: Auto<br>AGC: 300%<br>AcT: 60ms<br>CS: +2 to +5<br>IsM: Q<br>IsW: 1.6<br>Frag.: sHCD<br>NCE: 27, 32, 40 | classic orbitrap experiment: MS1 in Orbitrap at high resolution and data dependent MS2 in Iontrap at turbo scan rate. Dynamic exclusion enabled (exclude after n times=1; Exclusion duration (s)= 20; mass tolerance= $\pm 10$ppm).<br>Intensity Threshold: 25000 |
| ACE_0792 | Xcalibur<br>v4.5.445.18<br>Tune v3.5.3881.18<br>Gradient: 105 min | Analyzer: FT<br>Res.: 12000<br>SR: 375 - 1500<br>AGC: Standard<br>AcT: 50 ms<br>RF: 30<br>SF: --<br>DDM: CT/3sec | Analyzer: FT<br>Res./ScR: 15000/-<br>SR: Auto<br>AGC: 300<br>AcT: 60 ms<br>CS: +2 to +5<br>IsM: Q<br>IsW: 1.6<br>Frag.: sHCD<br>NCE: 27, 32, 40 | classic high high orbitrap experiment: MS1 in Orbitrap at high resolution and data dependent MS2 also in Orbitrap. Dynamic exclusion enabled (exclude after n times=1; Exclusion duration (s)= 20; mass tolerance= $\pm 10$ppm)<br>intensity threshold: $2.5 \cdot 10^4$ |

Note: **FT**= Fourier Transform (Orbitrap); **IT**= Iontrap; **Q**= Quadrupol; **Res.**= max. Resolution at 200 m/z (Lumos) or 400 m/z (Elite) [FWHM (full width at half maximum)]; **ScR**= scan rate for measurements in the IT; **SR**= scan range [m/z]; **AGC**= automatic gain control, max number of acquired ions per measurement; **AcT**= max. Ion acquisition time [ms]; **CS**= charge states used for fragmentation; **IsM**= Isolation mode (Q or IT), MS2 isolation and further is only done in IT; **IsW**= Isolation window [m/z], value followed by scan mode the isolation is based on (MS1, MS2 ...) **Frag.**= Fragmentation method; **HCD**= Higher-energy collisional dissociation; **CID**= Collision-induced dissociation; **ETD**= Electron-transfer dissociation; **EThcd**= Electron-Transfer/Higher-Energy Collision Dissociation; **sHCD**= stepped HCD; **NCE**= normalized collision energy; **cycles**: number of MSn recorded or max cycle time; **RF**= RF Lens [%]; **SF**= Source Fragmentation [V]; **DDM**: Data dependent Mode (cycle time in seconds, CT/[s] or number of scans, NS); **NS**= Number of data dependent scans.

## Notes

### Competing Interest Statement

The authors have declared no competing interest.

### Summary of Updates

This revised manuscript contains a much deeper analysis of the differental labeling sites triggered by ATP treatment.

